# Does post-ejaculatory sperm experience prior to fertilization alter offspring performance beyond paternity in an external fertilizing fish?

**DOI:** 10.1101/2024.04.18.590084

**Authors:** Ranjan Wagle, Craig F. Purchase

## Abstract

Recent research has suggested that the environment encountered by sperm post-ejaculation may impact offspring development beyond the transfer of the paternal genome. The mechanisms that underlie such effects remain unclear, but two non-mutually exclusive processes have been proposed. 1) Haploid selection, whereby stressful conditions act as stronger post-ejaculation pre-fertilization selective pressures on semen than under benign situations, resulting in the fertilization of, on average higher quality sperm and the production of offspring that exhibit superior performance across all environmental conditions that they might encounter. 2) Epigenetic inheritance, where environmental conditions induce changes in sperm that are passed down to offspring, resulting in altered gene expression in offspring. This would be adaptive if sperm experiences anticipate what is to come and improve offspring performance to match those conditions. Capelin (*Mallotus villosus*) sperm and embryos are sensitive to salinity and may represent a good system to investigate these phenomena. We used a split-ejaculate and split-brood experimental block design to expose capelin sperm to known benign (25 psu) and stressful (35 psu) salinity prior to egg contact, and split each batch of fertilized eggs for incubation at matched and mismatched salinity to those of sperm exposure. Our findings revealed no differences in hatch characteristics between offspring produced by sperm exposed to benign and stressful salinity conditions. A follow-up experiment found the same result with an increased selection gradient at 5 psu and 35 psu. Our study does not support the hypothesis that sperm experiences exert an influence on the development of offspring characteristics, independent of paternity. Instead, our results suggest that the sole influential factor in sperm determining offspring characteristics is the transfer of the paternal genome.

## Introduction

Offspring experience variable and potentially unpredictable conditions during their ontogeny. For species with external development this includes the embryonic stage, with high mortality and selection rates. When environmental conditions vary in space and across generations, selection encounters challenges in fine-tuning local adaptation (Mérot, 2022). This is complicated by a biphasic life cycle where selection pressures experienced by diploid adults may be very different from that of their haploid gametes (Purchase et al., 2021). Beyond direct genetic inheritance, parents possess the capacity to influence their offspring’s phenotype through other means, a phenomenon referred to as a parental effect (Badyaev & Uller, 2009). If these effects are adaptive, they can help offspring in coping to the actual conditions they encounter (Burgess & Marshall, 2014; Chirgwin et al., 2021; Crean et al., 2013; Jensen et al., 2014). Among parental effects, maternal effects hold particular significance in anisogamous reproduction, where the egg, characterized by its relatively large size and nutrient-rich composition, plays a pivotal role in embryo development. Compared to sperm, the substantial allocation of resources to eggs leads to a more direct and influential impact on the offspring’s development, particularly during critical early life stages (Bonduriansky & Head, 2007; Jensen et al., 2014; Mousseau & Fox, 1998; Wolf & Wade, 2009).

However, there is increasing evidence of paternal effects, challenging the conventional notion that fathers solely contribute DNA through sperm (Crean et al., 2013; Crean & Bonduriansky, 2014; Evans et al., 2017; Ragsdale et al., 2022; Simmons et al., 2022). Changes in the father’s experience, such as modifications in diet, changes in hormonal profiles, or altered social interactions, can affect the content of their semen and influence offspring development (Butzge et al., 2021; see review Evans et al., 2019; Ragsdale et al., 2022; Simmons et al., 2022). But sperm also encounter various conditions post-release, especially in external fertilizers (Purchase et al., 2021). Post-release sperm experiences influence the outcome of sperm competition and thus paternity, but additionally several studies have proposed that post-release sperm experiences may influence the subsequent phenotype of offspring outside of paternity (see review Crean & Immler, 2021). The literature presents contrasting perspectives on these effects. Some propose that preparing offspring for anticipated conditions could be an adaptive strategy (Immler et al., 2014; Ritchie & Marshall, 2013), while others express concerns about the potential transmission of physiological stress from sperm to offspring (Kekäläinen et al., 2018; Lymbery et al., 2021).

Sperm, beyond their role in transmitting haploid DNA, are equipped with a suit of epigenetic components such as DNA methylation, RNAs, and histone modifications (Immler, 2018). These components can undergo alterations in response to the post-ejaculatory environment and are integral to the process of epigenetic regulation, which affects gene expression without changing the DNA sequence (Immler, 2018; Lettieri et al., 2019b; Lymbery et al., 2020; Pitnick et al., 2020). However, for epigenetics to play an adaptive role, these altered sperm must impact the phenotype of the resulting offspring (Donkin & Barrès, 2018; Marshall, 2015). These alterations in sperm may confer advantages to offspring when they encounter similar conditions to those experienced by the sperm (Figures 1 A & B). This introduces the anticipatory hypothesis, suggesting that offspring are primed for anticipated conditions, enhancing their performance in such conditions (Burgess & Marshall, 2014; Graziano et al., 2023; Hosken et al., 2003; Marshall, 2015; Ritchie & Marshall, 2013). However, if conditions differ from those experienced by the sperm, offspring performance may not improve, as the modifications may not prove advantageous in the new context (Lymbery et al., 2021).

**Figure 1.**
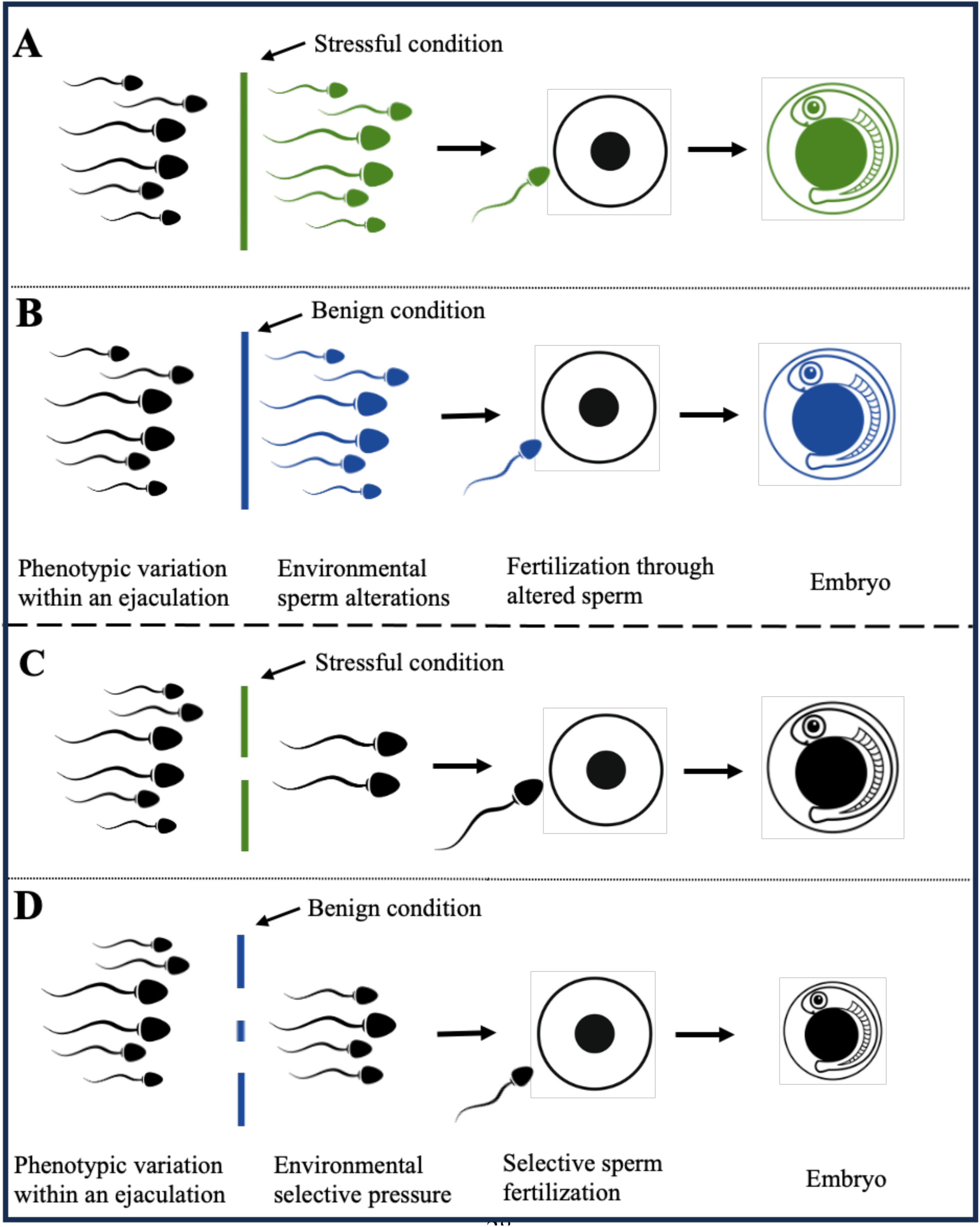
Impact on embryo development resulting from sperm experiences through two potential non-mutually exclusive mechanisms: epigenetic alterations (panels A and B) and haploid selection (panels C and D). In panels A and B, stressful condition (A) and benign condition (B) induce alteration in sperm (indicated by colour), affecting all sperm. These altered sperm influence embryo performance in such a way that embryos show optimal performance when exposed to condition matching to those of sperm exposure (same colour of sperm and embryo), as opposed to mismatched condition. This effect on the embryo due to the alteration of sperm can also be called as modifying/anticipatory effect. Whereas, in panels C and D, under stressful condition (C), a higher selection pressure acts upon sperm, leading to the preferential selection of, on average, higher-quality sperm. These selected higher-quality sperm contribute to superior embryo performance across all conditions (as indicated by embryo size), whereas in benign condition (D), lower selection pressure permits, on average, lower-quality sperm pass to through, resulting in embryos of reduced overall quality. This effect on the embryo due to the selection of sperm can also be called a filtering effect.

In contrast to sperm modifications inducing parental effects in offspring, there is also the potential possibility for haploid selection (Immler et al., 2014; Kekäläinen et al., 2018; Lymbery et al., 2021; Ritchie & Marshall, 2013). Sperm competition is usually conceptualized as between ejaculates of rival males (Parker, 2020). However, the presence of numerous sperm cells in an ejaculate, along with the limited availability of eggs for fertilization, naturally gives rise to competition among sperm within an ejaculate (Immler et al., 2014; Sutter & Immler, 2020). This competition is significantly influenced by the phenotypic traits of individual sperm, which include morphology, motility, and viability (Immler et al., 2008). Traditionally, these traits were thought to be regulated only by the male’s diploid genotype (all sperm within an ejaculate, thus having phenotypes under the same genetic influence), hence the view that each sperm is a mere carrier of unique paternal genetic information without influence on sperm phenotypes. However, emerging studies may suggest that these phenotypic traits are partly genetically determined by the sperm’s haploid DNA (Alavioon et al., 2017; Borowsky et al., 2018; see review Immler, 2019). This introduces the concept of haploid selection, a process whereby the phenotype of sperm is partially governed by their unique haploid genome (Alavioon et al., 2017; Immler et al., 2014). If this is true, sperm with “high quality” genes can manifest phenotypically superior sperm traits (for example, larger size, higher motility) and may confer a selective advantage that can impact fertilization outcomes.

Conditions faced by sperm post-release further influence the outcome of this competition among sperm within an ejaculate. Post-release under benign conditions, where the selection pressure is lower, a greater number of lower quality sperm can achieve fertilization, lowering the average quality across embryos. This, in turn, can lead to the production of offspring of average lower quality. In contrast, under stressful conditions, due to higher selection pressure, on average, superior sperm are selected (Holt & Look, 2004; Sutter & Immler, 2020). This selective filtration can result in a greater proportion of the remaining superior sperm carrying ’high-quality’ genes than under benign conditions. Consequently, this may result in the production of, on average, high-quality offspring capable of superior performance across diverse environmental conditions (Figures 1 C & D).

Therefore, beyond paternity (among males), offspring performance is possibly influenced by sperm experience (Purchase et al., 2021), encompassing which sperm within an ejaculate fertilizes the egg (haploid selection) and what happened to the fertilizing sperm (epigenetics). These mechanisms collectively contribute to shaping the genetic and epigenetic inheritance that offspring receive from their fathers, ultimately influencing both offspring genotype and phenotype. Therefore, it is imperative to conduct further research to understand these intricacies and ascertain their relative contributions. Such understanding is important in elucidating the potential adaptive mechanisms at play and thereby shedding light on evolutionary processes and local adaptation.

To explore which potential mechanisms, haploid selection or epigenetics, have a greater influence on offspring, we used the marine beach spawning fish, capelin (*Mallotus villosus*), as its sperm and embryos are highly sensitive to salinity (Beirão et al., 2018a; Purchase, 2018). Capelin have a unique reproductive adaptation/constraint for an external fertilizer wherein its sperm are active upon leaving the male’s body, and subsequent sperm swimming is affected by salinity (Beirão et al., 2015; Beirão et al., 2018a). Capelin embryos, although sensitive to oceanic salinity, do develop successfully in nature (Purchase, 2018). This could be due to the potential transmission of adaptive traits regarding salinity conditions from the post-ejaculation experiences of capelin sperm to embryos, possibly enhancing the embryo’s ability to cope under such conditions. To investigate the existence of such an effect of capelin sperm and, if confirmed, to delineate the underlying mechanisms and their relative contribution, we exposed sperm to benign or stressful salinity conditions and then incubated the embryos under conditions that either corresponded to or differed from the sperm’s initial salinity exposure. Specifically, if there are no differences between offspring development sired by sperm exposed to different conditions, it implies there is no impact of sperm experiences on offspring development (Figure 2 A). If we observe improved offspring development when the incubation conditions match those experienced by the sperm, it would point toward the inheritance of epigenetic modifications (Figure 2 B). Conversely, if we find that offspring development is superior when sired by sperm exposed to stressful salinity conditions, regardless of the salinity conditions during embryo incubation, it suggests the haploid selection process (Figure 2 C).

**Figure 2.**
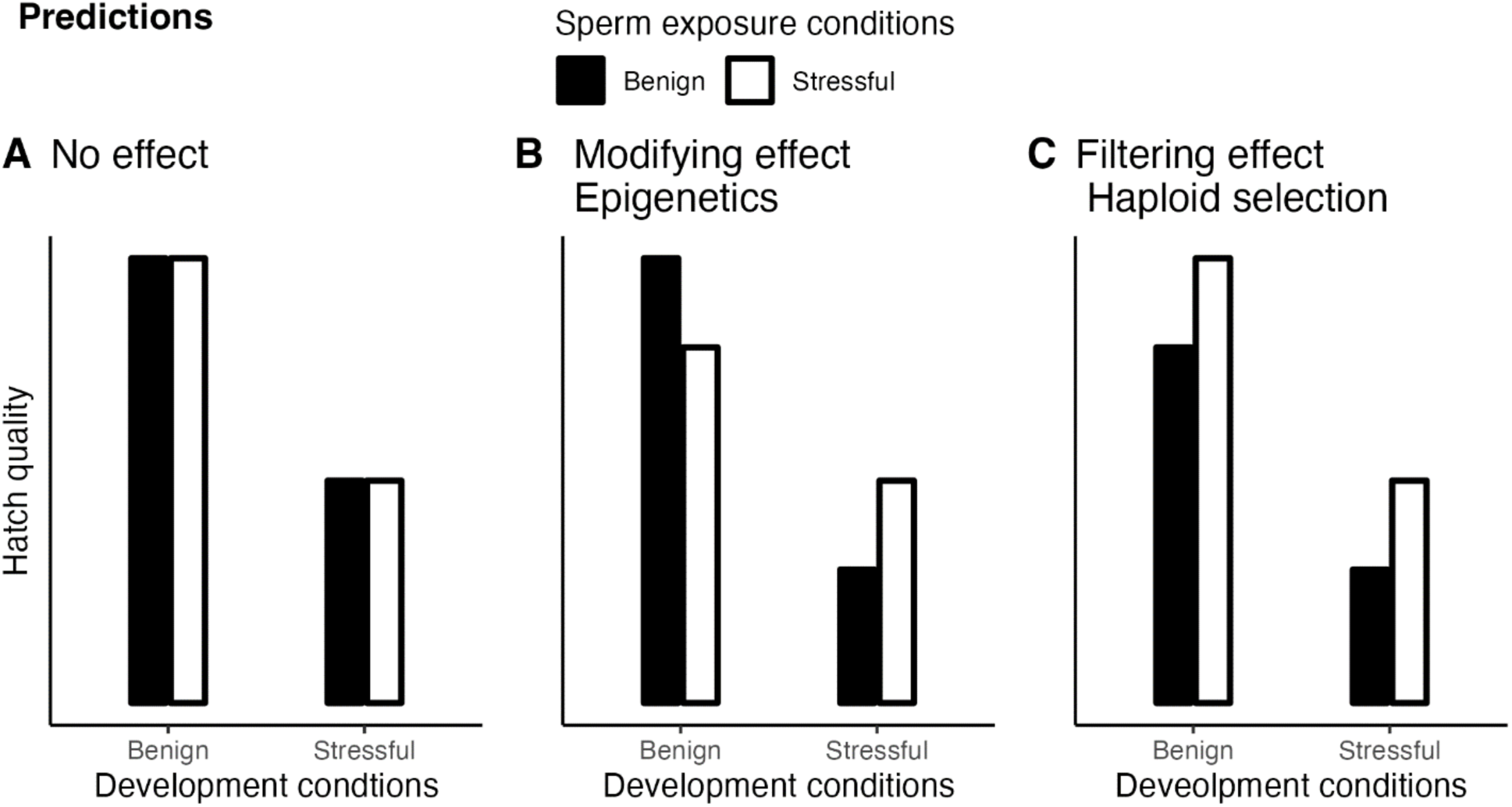
Predictions on the effect of pre-fertilization sperm exposure to benign and stressful conditions on hatch quality under matched and mismatched incubation conditions to those of sperm exposure. Shown are three panels representing different possible outcomes: (A) No effect, where exposure of sperm to different conditions has no impact on hatch quality; (B) Modifying effect, where exposure of sperm induces modifications within the sperm themselves, resulting in better hatch quality under matched versus mismatched conditions to those of sperm exposure and (C) Filtering effect, where exposure of sperm to stressful conditions filters out more of the low-quality sperm, resulting in better hatch quality in all conditions due to, on average, to fertilization from sperm of higher quality.

## Methodology

### General design

The study used a split-ejaculate and split-brood experimental block design to investigate the influence of sperm exposure on offspring development (Figure 3). To isolate the influence of sperm post-ejaculation pre-fertilization exposure on offspring, offspring performance sired by sperm exposed to benign or stressful conditions was compared under matched or mismatched conditions to those of sperm exposure. In 2022, 33 experimental blocks (families) were used with a “lower” exposure gradient, while in 2023, a “higher” exposure gradient was implemented, employing 9 blocks (families). Each block was based on a different male-female pairing to control for any systematic variation due to diploid paternal and maternal level effects.

**Figure 3.**
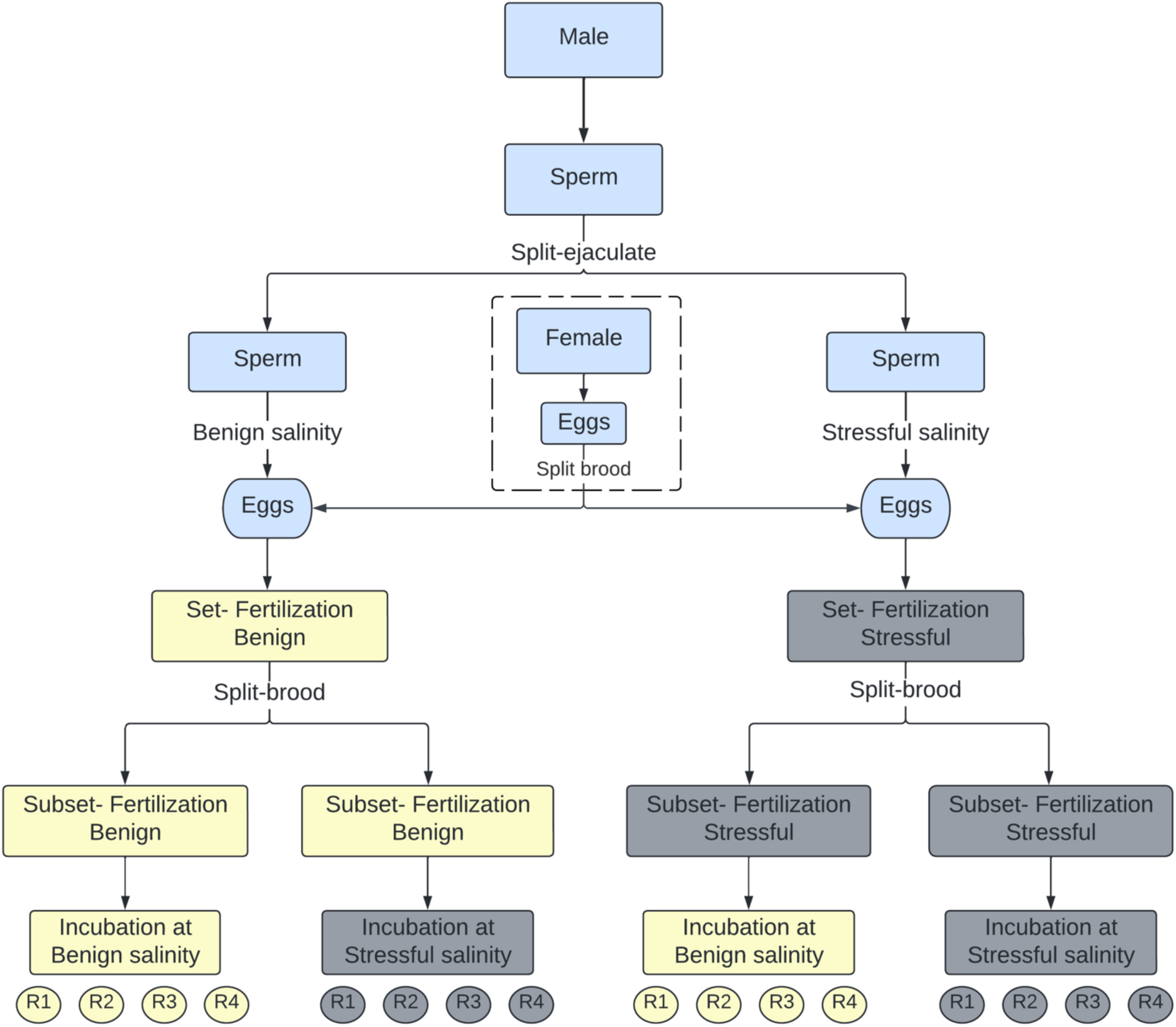
Split-ejaculate and split-brood experimental design to investigate the effect of capelin pre-fertilization sperm exposure to two salinities on offspring development at matched and mismatched salinity level to those of sperm exposure. Shown is the procedure for 1 family (block) created from 1 male and 1 female chosen at random. Eggs from the single female were split into two standardized batches. A male’s ejaculate was then divided into two standardized aliquots, with one exposed to benign salinity and the other to stressful salinity for 4 seconds prior to egg contact. Each brood of fertilized eggs was further split into two subsets and incubated at matched and mismatched salinity to those of sperm exposure. The design controlled for male and female variation and allowed the isolation of the effect of sperm exposure to different salinities on offspring development. R1, R2, R3, and R4 are replicate beakers. 2022 experiment - Lower exposure gradient: Benign (25 psu), Stressful (35psu), n = 33 blocks/families 2023 experiment - Higher exposure gradient: More benign (5 psu), Stressful (35psu), n = 9 blocks /families

### 2022 experiment: lower exposure gradient

#### Capelin collection and gamete collection

Capelin were captured using a cast net in July 2022 as they were spawning on four beaches of Newfoundland, Canada (Middle Cove: 47°39’ 2.75" N, 52°41’ 45.24" W; Bellevue Beach: 47° 38’ 9.276" N, 53° 46’ 51.0924"W; Bryant’s Cove 47° 34’ 18.155" N, 52° 44’ 14.862" W; Chapel Arm: 47° 31’ 0.6564" N, 53° 40’ 10.7508" W). They were transferred to the laboratory and kept in flow-through (simulated natural photoperiod) seawater tanks (males and females separately) at 4 – 8 °C that roughly matched the conditions in the natural habitat at the spawning period. Gametes were collected within 48 hours of capture.

To create an experimental block, a randomly selected female capelin was euthanized using an overdose of tricaine methanesulfonate (also known as MS-222) buffered with sodium bicarbonate and then dried using a paper towel. The eggs were collected through gentle pressure on the abdomen (Beirão et al., 2018a; Purchase, 2018), divided into two standard batches of 0.4g each and placed in flexible teflon trays on ice. Similarly, semen from a male was collected and split into two 20 μL aliquots. Only males with whiter semen, indicative of high sperm density (Steyn, 1993), were used to ensure good fertilization success in the experiment (the goal was to monitor offspring characteristics).

#### Sperm exposure

Capelin sperm are motile upon release from the male body and exhibit salinity sensitivity (Beirão et al., 2018a). Hence, without delay, to manipulate the environment that semen (hereafter sperm) experienced, each aliquot was exposed to 40 μL water of either benign (25 psu) or stressful (35 psu) salinity (based on Beirão et al., 2018a; Purchase, 2018) for a period of 4 seconds (Figure 3), then the sample was returned to a salinity of 30 psu (midpoint of 25 psu and 35 psu) by exposing it to reciprocal salinity for 1 second. The purpose of returning the sample to 30 psu was to ensure that the eggs received the same salinity level regardless of which salinity exposure the sperm had undergone. This ensured that any effects on the embryos were solely due to sperm exposure.

The exposure gradient, exposure duration, and sperm-water ratio were chosen to optimize their impact on embryo development while carefully considering the trade-off involved in maintaining a sufficient number of embryos produced (fertilization success could not be too low). Additionally, it should reflect realistic natural conditions. Specifically, 25 psu and 35 psu were chosen as 35 psu is stressful to capelin sperm and embryos, and slightly higher than typical salinity levels found in costal seawater, while 25 psu, is more benign (Beirão et al., 2018a; Purchase, 2018). Capelin sperm are already active upon release, so 4 seconds of exposure duration was selected to optimize the effect on embryos while still yielding enough fertilized eggs for measuring various hatch characteristics, recognizing its potential impact on fertilization success. Additionally, a sperm-water ratio of 1:2 was chosen to dilute the semen enough to dramatically change the chemistry of the seminal plasma while ensuring an adequate number of fertilized eggs. Salinity water used for sperm exposure was maintained at 4 °C.

#### Invitro-wet fertilization

Immediately after sperm exposure, within each block, to create two sets of fertilized eggs of benign and stressful salinity, each exposed sperm aliquot was mixed with a standard batch of eggs (i.e. 0.4g of eggs) using a toothpick. Care was given to ensure that both standard batches of eggs underwent the same procedure and fertilization was done at the same time. Capelin eggs are sticky when they come in contact with water, so immediately after fertilization, a solution of 600 mg/L tannic acid (see Purchase (2018)) mixed in 30 psu water was added and swirled for 30 seconds to remove stickiness. This was then poured off and rinsed three times with 30 psu water, making the observation of individual eggs possible.

For each block, two control batches underwent the same process as the fertilized eggs described above but did not have sperm added. These unfertilized eggs were used as a reference when assessing fertilization success.

#### Incubation

The two sets of fertilized eggs per block were divided into four fertilization subsets based on two sperm salinity × incubation salinity combinations, each consisting of approximately 1200 eggs. The subsets were then subjected to incubation at either matched or mismatched salinity to that of sperm exposure, resulting in four distinct conditions: benign-benign, benign-stressful, stressful-benign, and stressful-stressful (see Figure 3). Additionally, the two control batches per block were incubated, one at benign and the other at stressful salinity. All four fertilization subsets and two control batches were incubated at 4 °C.

After 20 hours, approximately 100 eggs from each fertilization subset were examined under a microscope to determine fertilization success by counting the number of fertilized (8 and 16-cell differentiation stages) and unfertilized eggs (no cell division) (Beirão et al., 2018b). To optimize our efforts in ensuring successful hatches to produce enough data for the analysis of offspring performance, families with > 25% average fertilization success (n=33 in 2022) were chosen to proceed with the experiment. The fertilization rates of two families were assessed twice using different eggs within each subset to measure the repeatability of our assessment (Wagle, 2024).

Following previous protocols to transfer the eggs to replicate incubation beakers (Purchase, 2018), a small group of eggs was removed from a fertilization subset, placed in a petri dish and counted. Additional eggs were added or removed until the final count reached 50 eggs. A picture was taken to validate the count. The eggs were then transferred to a replicate beaker containing 40 mL of water of the same salinity as that of the subset. This process was repeated until each family was transferred to 16 beakers, with four replicate beakers for each subset (4 subsets × 4 replicate incubation beakers = 16 beakers).

These replicate incubation beakers were kept in the same incubator, and the temperature was raised from 4 °C to 10 °C. This temperature was chosen to produce the highest hatch success rates across all salinity levels (Purchase, 2018). This choice is important because it ensured a higher sample size to measure other offspring development parameters, which rely on hatch success. All replicate beakers from the same family were placed in the same incubator. The water in each replicate beaker was partially replaced with new water of the same salinity every other day, with half of the volume being decanted off before the addition of new water. Experimental saline water was prepared in bulk by dissolving an appropriate amount of Instant Ocean Sea Salt^TM^ in dechlorinated tap water and stored at 10 °C.

Replicate beakers were checked daily at 10 AM for hatch larvae, and the larvae were removed immediately and recorded by replicate beakers. All larvae were preserved in buffered formalin solution (2.2%), and hatch size was determined using digital pictures taken from a dissecting microscope. For a given replicate incubation beaker, once hatching started, if no new larvae hatched for three consecutive days, it was assumed that no more would hatch, and the replicate beaker was discarded (Purchase, 2018).

### 2023 experiment: higher exposure gradient

In 2022, we found no significant effect of sperm exposure on offspring traits (see result section). So, in 2023, we increased the exposure gradient from the previous experiment in the hope of gaining more insights into the impact of sperm exposure on offspring development and measured the length of the starvation time to assess the potential effects of sperm exposure on offspring in later stages of life (allowing more time for the paternal genome to have an effect).

The fertilization and incubation process followed the protocol of the 2022 experiment, with some modifications. Gametes were collected from fish within 6 hours of capture. To increase the exposure gradient (Purchase 2018), sperm aliquots were exposed to 5 psu (benign) and 35 psu (stressful) at a sperm-to-salinity water ratio of 1:5. The tannic acid solution was prepared using 20 psu water (midpoint of 5 psu and 35 psu). To improve the assessment of fertilization success, this time, approximately 300 eggs were scored. Consistency in our scoring was evaluated by scoring two sets of eggs twice of two families (Wagle, 2024), resulting in increased scoring precision compared to 2022.

The initial five larvae hatched from each beaker were preserved in a 2.2% buffered formalin solution for hatch size measurement. Subsequently, to measure the length of starvation time, a minimum of ten larvae were collected from each beaker carefully using a wide-mouth disposable pipette, either on the same day or spread across subsequent days, based on availability (Purchase, 2018). These larvae were then transferred to 50 mL glass vials filled with the same salinity water as the beaker from which they were collected and kept at 10 °C. The number of days required for all larvae to die from the day of hatch was recorded as starvation time. Each vial held a maximum of five larvae, and new vials were used daily. Larvae were checked daily, and survivors were counted and recorded (Purchase, 2018). The remaining hatched larvae were preserved in the same formalin solution along with the initial first five larvae. From these preserved larvae, five larvae were randomly selected for hatch size measurements.

### Calculations and statistical analysis

The calculation of hatch time involved computing a weighted average by multiplying the number of larvae hatched on a given day by the number of days it took them to hatch, summing the values across all hatching days, and dividing by the total number of hatched eggs (Purchase, 2018). As we had four values from four replicate beakers for all hatch characteristics for each salinity treatment (except in some treatments in hatch time, hatch size and starvation time), a data quality control approach was taken where any experimentally produced outliers were identified using a coefficient of variation above 30% to address possible measurement errors (Brown, 1998). Outliers identified were excluded, and the remaining data were used for analysis. Outliers were only observed in the hatch success data, with no outliers detected in the datasets for hatch time and hatch size in both 2022 and 2023. This approach was not extended to the starvation time data due to limited data availability.

Hatch characteristics of four replicate beakers, if available, containing the same salinity water from each fertilization subset were averaged to generate one value (per fertilization subset) and used in data analysis (Figure 3). Starvation time was determined only in 2023; the values were averaged first between vials (if available) and then between four replicate beakers. The data for hatch and starvation time had a resolution of a whole day.

Each hatch characteristic was analyzed separately for each year, as treatment levels varied between years. Odds ratios of events (number hatched) and trails (total number of eggs per beaker) were analyzed using a generalized linear mixed model with binomial error distribution. The model included fixed-effect categorical variables of sperm salinity, incubation salinity, and their interaction, as well as a random effect of family. Additionally, only in 2023, fertilization success was added as a covariate to the model and reanalyzed. Due to low consistency in scoring fertilization success in 2022, this step was not taken during that year’s analysis. The same model parameters were used for hatch time, starvation time and hatch size analyses but utilizing a general linear mixed model with a normal error distribution. Statistical analyses were performed in R v. 4.3.0 (R Core Team, 2023) using the ‘lme4’ package (Bates et al., 2015).

## Results

If offspring development improves when the incubation conditions match those experienced by the sperm, it would support the epigenetic hypothesis with a significant interaction between sperm salinity and incubation salinity. Our results (Tables 1 & 2) indicate no significant interactions on all hatch characteristics in both 2022 and 2023. If offspring sired by sperm exposed to stressful salinity condition exhibit higher development across all salinity conditions, then it would support the haploid selection hypothesis with significance of sperm salinity in the model. We found that sperm exposure did not influence development at either offspring salinity (Tables 1 & 2). As expected, there was a positive relationship between fertilization rate and hatch success (Table 1, 2023a). We observed that benign incubation salinity conditions led to higher hatch success (∼1.12 times), shorter hatch time (∼1.02 times) and hatch at a larger size (∼1.03 times) in 2022 when compared to the stressful incubation salinity condition. Notably, this contrast was more pronounced in 2023 (∼4.5 times for hatch success calculated from both total and fertilized eggs, 1.13 times for hatch time and 1.13 for hatch size) than in 2022 (Figures 4 A-D and Figures 5 A-C). Starvation time was only measured in 2023 and showed the same pattern: ∼2.19 times longer time to starve in benign incubation salinity than in stressful incubation salinity (Figure 4 E) but non-significant.

**Figure 4.**
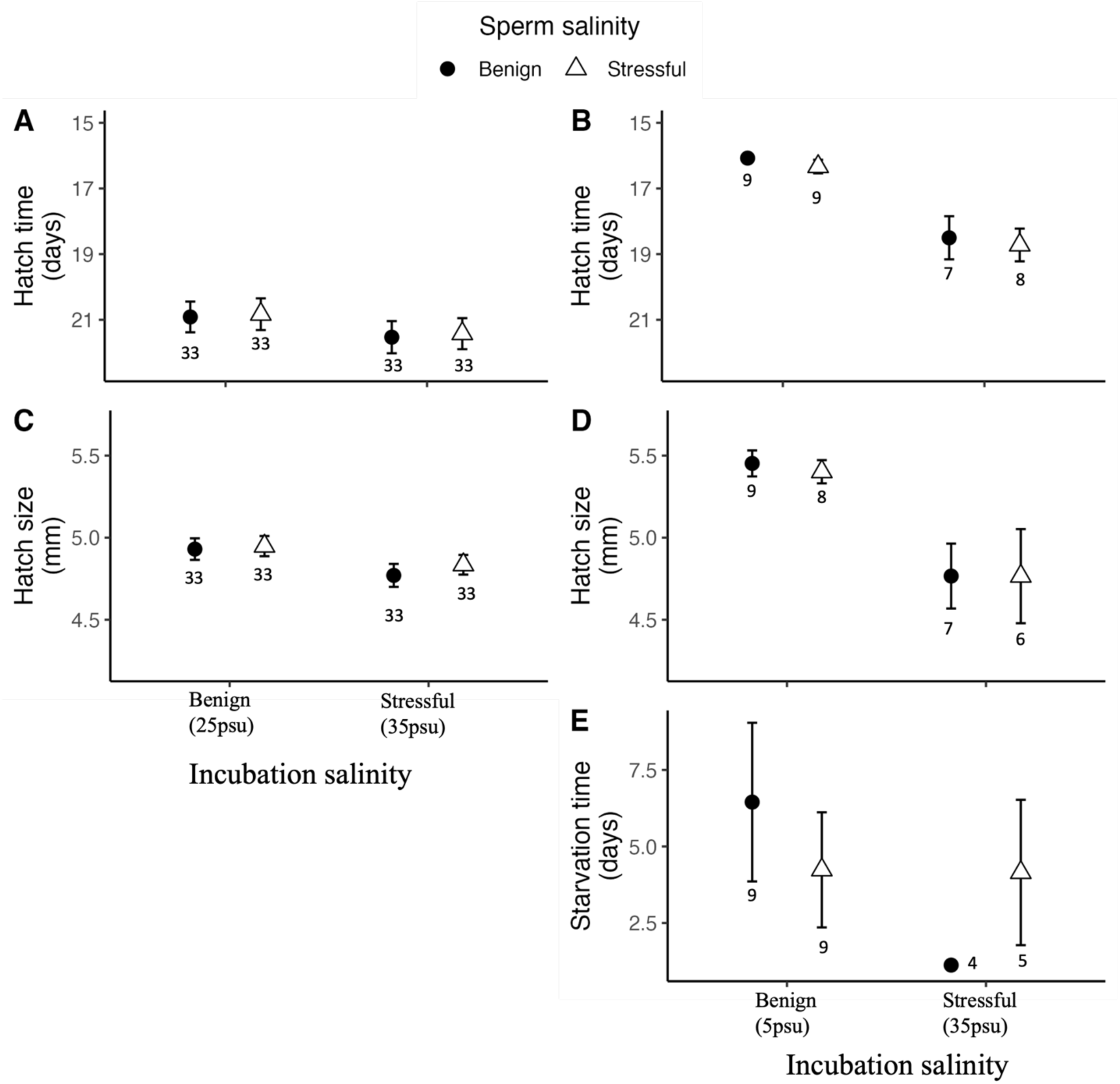
The effects of sperm and incubation salinity using a split-ejaculate and split-brood experimental design on hatch qualities of capelin. The left panel presents hatch time (A) and hatch size (C) from 2022, while the right is hatch time (B), hatch size (D), and starvation time (E) from 2023. Data are shown as means, averaged within families and treatments, calculated giving equal weight to each replicate incubation beaker. For starvation time, equal weighting was first done within vials and then across incubation beakers, treatments, and families. Error bars are standard errors among families, with the number next to each point indicating the number of families. Family number differences in hatch characteristics, apart from hatch success, in some treatments in 2023 are due to no hatching in those treatments. Y-axis of hatch time is inverse, indicative of its negative association with hatch quality. Salinity conditions in 2022 were benign (25 psu) and stressful (35 psu), while in 2023, benign at 5 psu and stressful at 35 psu (Purchase, 2018).

**Figure 5.**
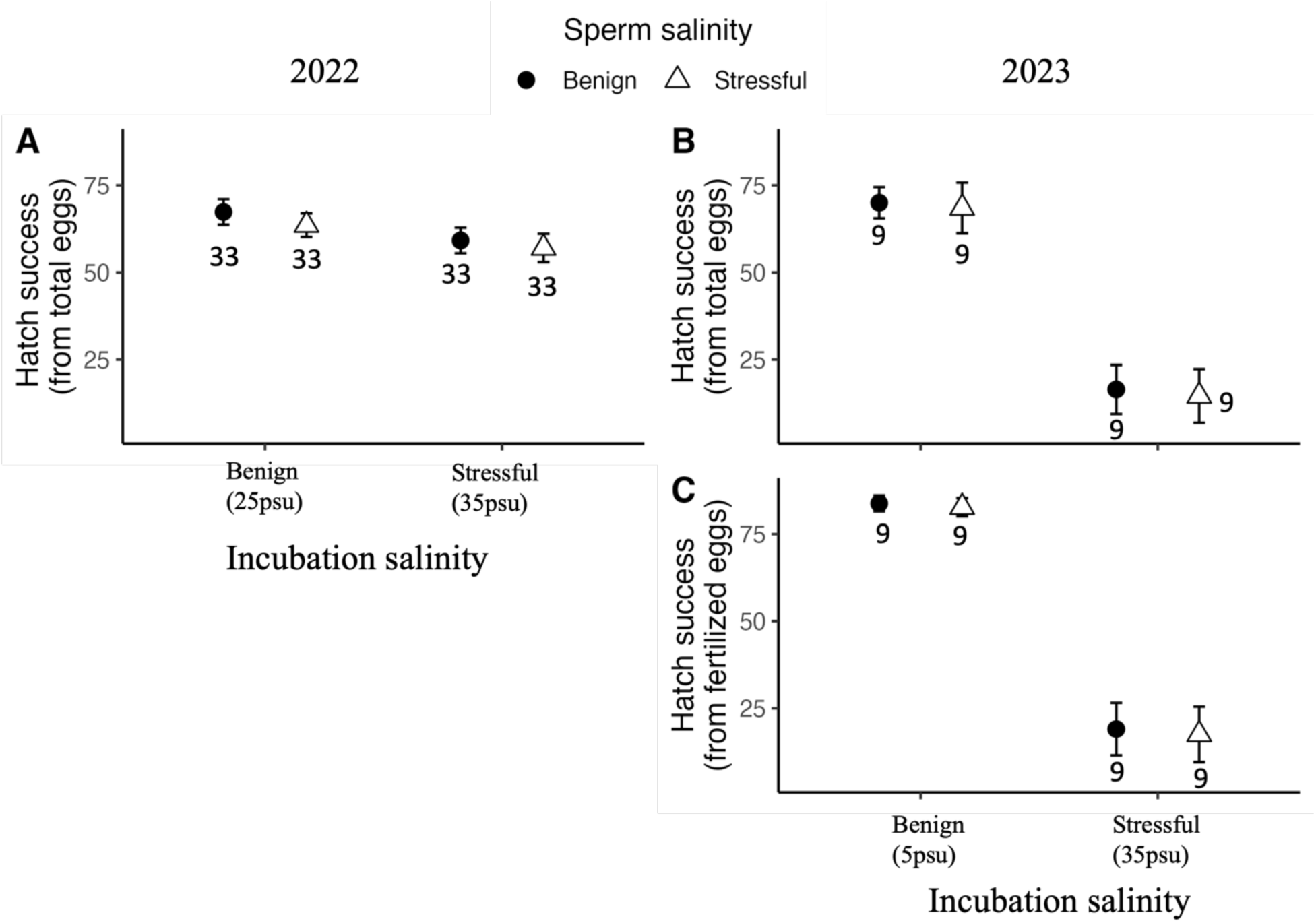
The effects of sperm and incubation salinity using a split-ejaculate and split-brood experimental design on hatch success. The left panel presents hatch success, calculated from total eggs (A) from 2022, while the right is hatch success, calculated from total eggs (B), and hatch success, calculated from fertilized eggs (C) from 2023. Data are shown as means, averaged within families and treatments, calculated by giving equal weight to each replicate incubation beaker. Error bars are standard errors among families, with the number next to each point indicating the family numbers. Salinity conditions in 2022 were benign (25 psu) and stressful (35 psu), while in 2023, benign at 5 psu and stressful at 35 psu (Purchase, 2018).

**Table 1.** Type III Wald χ2 tests for the effects of sperm exposure salinity, incubation salinity and their interaction (S × I) on odds of hatching in capelin, fitted using generalized linear mixed effects models (GLLM), in 2022 and 2023 (without fertilization covariate) and 2023a (with fertilization covariate). Removing the interaction term from the models did not change the result of odds of hatching in either year (results not shown). The results also remained consistent when re-running the models with the data without removing outliers, with no change observed upon excluding the interaction term (results not shown). * Indicates p-value < 0.05.

| Effects | 2022 |  |  | 2023 |  |  | 2023a |  |  |
| --- | --- | --- | --- | --- | --- | --- | --- | --- | --- |
| | $\chi^2$ | d.f. | p-value | $\chi^2$ | d.f. | p-value | $\chi^2$ | d.f. | p-value |
| Intercept | 29.102 | 1 | < 0.001* | 7.269 | 1 | 0.007* | 24.654 | 1 | < 0.001* |
| Sperm salinity | 0.949 | 1 | 0.329 | 0.088 | 1 | 0.766 | 0.059 | 1 | 0.806 |
| Incubation salinity | 5.203 | 1 | 0.023* | 113.24 | 1 | < 0.001* | 106.76 | 1 | < 0.001* |
| $S \times I$ | 0.011 | 1 | 0.917 | 0.017 | 1 | 0.8952 | 0.016 | 1 | 0.896 |
| Fertilization (covariate) |  |  |  |  |  |  | 17.755 | 1 | < 0.001* |

**Table 2.** Type III Analysis of variance table with Satterthwaite’s method for the effects of sperm exposure salinity, incubation salinity and their interaction (S × I) in 2022 on a) hatch time, b) hatch size in capelin. Additionally, in 2023, the analysis included c) starvation time. Linear mixed effect models (LLM) fit by residual maximum likelihood (REML) in both 2022 and 2023. Removing the interaction term from the models did not change the results for any hatch characteristics in either year (results not shown). * Indicates p-value < 0.05.

| Effects | 2022 |  |  | 2023 |  |  |
| --- | --- | --- | --- | --- | --- | --- |
|  | F-value | d.f. | p-value | F-value | d.f. | p-value |
| a) Hatch time |  |  |  |  |  |  |
| Sperm salinity | 2.175 | 1, 96 | 0.144 | 0.442 | 1, 21.82 | 0.512 |
| Incubation salinity | 87.790 | 1, 96 | $< 0.001^*$ | 48.923 | 1, 22.08 | $< 0.001^*$ |
| $S \times I$ | 0.039 | 1, 96 | 0.845 | 0.0084 | 1, 22.08 | 0.927 |
| b) Hatch size |  |  |  |  |  |  |
| Sperm salinity | 1.271 | 1, 96 | 0.262 | 0.038 | 1, 21.26 | 0.847 |
| Incubation salinity | 13.899 | 1, 96 | $< 0.001^*$ | 18.692 | 1, 18.20 | $< 0.001^*$ |
| $S \times I$ | 0.420 | 1, 96 | 0.528 | 0.007 | 1, 18.43 | 0.935 |
| c) Starvation time |  |  |  |  |  |  |
| Sperm salinity |  |  |  | 0.103 | 1, 16.34 | 0.752 |
| Incubation salinity |  |  |  | 1.640 | 1, 17.85 | 0.216 |
| $S \times I$ | | | | 1.827 | 1, 16.34 | 0.194 |

## Discussion

Juveniles may encounter unpredictable environmental conditions during development due to spatial or temporal variation in the landscape or environmental stochasticity across generations. This creates a challenge for selection to fine-tune local adaptation in development, especially in externally fertilizing species where high embryo mortality results in higher selection pressures. Parental effects may bridge this gap by transmitting information to offspring about conditions they are likely to encounter, which can influence their development. In external fertilizers, prior to fertilization, sperm are exposed to conditions that embryos will likely experience. Sperm experiences may have the potential to alter embryo development in an adaptive way. We tested this in beach spawning capelin, where development occurs across widely varying salinities. Capelin sperm swimming performance and embryo development are both highly sensitive to external salinity, however, our results indicate that sperm exposure to benign or stressful salinity conditions does not influence offspring development (i.e. hatch time and size, and starvation time). This suggests that, contrary to some previous studies in other species, sperm experiences do not serve as a conduit for parental effects in capelin; seemingly, a sperm’s only role is transferring the paternal genome. But further research is required to clarify whether sperm experiences, beyond salinity exposure, exert an influence on offspring development in capelin.

The influence of sperm post-ejaculation experiences on offspring phenotype is a subject of ongoing research yielding a range of results. Lymbery et al. (2021) revealed that exposure of sperm to high temperatures, indicative of stressful conditions, exerted adaptive effects on embryos of mussel (*Mytilus galloprovincialis*) only when the embryos were incubated at ambient temperatures, considered benign conditions. However, when embryos were incubated at high temperatures, those sired by sperm treated at high temperatures exhibited inferior performance. In contrast, Ritchie and Marshall (2015) and Graziano et al. (2023) suggested that the adaptive effects of tubeworm (*Galeolaria gemineoa*) and salmon (*Salmo salar*) sperm exposure when the offspring condition aligns with the condition of sperm exposure, hinting at the epigenetic mechanisms at play. On the contrary, Kekäläinen et al. (2018) present evidence of maladaptive effects on European whitefish (*Coregonus lavaretus*) offspring resulting from sperm experiences, further complicating the overall picture. It is important to note that our study represents a departure from these findings, as it reports no discernible impact of sperm experiences on the offspring phenotype.

We propose two primary interpretations for our findings. Firstly, it is well-established that sperm possess a group of epigenetic components (Donkin & Barrès, 2018; Immler, 2018) and undergo changes due to exposure to post-release conditions (Lymbery et al., 2020; Pitnick et al., 2020). But, for any post-release alterations in these epigenetic components to affect the embryo, they must have a functional role during the embryo’s development. Our study suggests that salinity exposure either does not alter the epigenetic component of capelin sperm or that if it does; the changes do not translate into developmental effects on the embryo. This implies that the post-release epigenetic condition of sperm does not affect the embryo, possibly because these changes do not persist through fertilization or are not influential during the critical stages of embryonic development. Secondly, the post-release condition may selectively influence the average phenotype of sperm that is able to fertilize eggs (Alavioon et al., 2017; Marshall, 2015). However, our results suggest no effect of sperm salinity experiences on embryo development; therefore, it appears that phenotypic variation among sperm within a single ejaculate may not be influenced by their haploid genome. Taken together, these findings align with the traditional belief that the function of sperm is predominantly to deliver the paternal genome and that the experiences of sperm do not impart developmental directives to the embryo, at least in capelin and in the context of salinity.

## Acknowledgements

Assistance in collecting capelin and conducting experiments was provided by Nick Murphy, Kaitlyn Gladney, Gage Moffatt, Maliya Cassels, Alyssa Forget, Johanna Bosch, Tatiana Hyde and Connor Hanley. Funding was provided via Memorial University and grants to Craig Purchase from the Natural Sciences and Engineering Research Council of Canada, the Canada Foundation for Innovation, and the Research and Development Corporation of Newfoundland and Labrador. All procedures followed the Canadian Council on Animal Care guidelines for the use of research animals (Memorial University protocols 18-02-CP, 22-02-CP) and Fisheries and Oceans Canada permits (NL-6666-22 and NL-7297-23).

